# Regeneration of a Full-Thickness Defect in Rotator Cuff Tendon with Umbilical Cord-Derived Mesenchymal Stem Cells in a Rat Model

**DOI:** 10.1101/2020.06.12.148544

**Authors:** Ji-Hye Yea, InJa Kim, Gayoung Sym, Jin-Kyung Park, Ah-Young Lee, Byeong Chan Cho, Tae Soo Bae, Byoung Jae Kim, Chris Hyunchul Jo

## Abstract

Although rotator cuff disease is a common cause of shoulder pain, there is still no treatment method that could halt or reveres its development and progression. The purpose of this study was to investigate the efficacy of umbilical cord-derived mesenchymal stem cells (UC MSCs) on the regeneration of a full-thickness rotator cuff defect (FTD) in a rat model. We injected either UC MSCs or saline to the FTD and investigated macroscopic, histological and biomechanical results and cell trafficking. Treatment with UC MSCs improved macroscopic appearance in terms of tendon thickness at two weeks, and inflammation, defect size, swelling/redness and connection surrounding tissue and slidability at four weeks compared to the saline group. Histologically, UC MSCs induced the tendon matrix formation recovering collagen organization, nuclear aspect ratio and orientation angle of fibroblast as well as suppressing cartilage-related glycosaminoglycan compared to saline group at four weeks. The UC MSCs group also improved ultimate failure load by 25.0% and 19.0% and ultimate stress by 27.3% and 26.8% at two and four weeks compared to saline group. UC MSCs labeled with PKH26 exhibited 5.3% survival at four weeks compared to three hours after injection. This study demonstrated that UC MSCs regenerated the FTD with tendon tissue similar properties to the normal tendon in terms of macroscopic, histological and biomechanical characteristics in a rat model.

## Introduction

Shoulder pain results in over 4.5 million physician visits, and rotator cuff disease is the most common cause of shoulder pain accounting up to 70% of cases with approximately 300,000 operations each year in the United States(1, 2). The spontaneous healing potential of tendon is very limited(3). Once injured or diseased, tendinopathy typically progresses without recovery. Current conservative treatments for rotator cuff disease include rest, nonsteroidal anti-inflammatory drugs, physical therapy and various kinds of injections(4). However, the success rate for conservative treatment varies widely from less than 50% to greater than 90%(5). A considerable number of patients, 41%, showed persistent symptoms after 12 months of treatment(6). These results may be attributed to the fact that those symptomatic treatments do not address the fundamental etiology of rotator cuff disease, most importantly tendon degeneration(7). As tenocytes in degenerative tendon do not participate in the regeneration process(8), it would be momentous to investigate new biological strategies such as application of various cells, growth factors and cytokines to more adequately treat tendon degeneration(9).

A promising therapeutic strategy for recovering damaged tissues is the use of mesenchymal stem cells (MSCs) which may be derived from various tissues such as bone marrow (BM), adipose tissue (AD), muscle tissue (MT) and umbilical cord blood (UCB)(10). Recently, a few studies have shown that MSCs were effective in regenerating rotator cuff tendon(11, 12). BM MSCs improved early collagen organization and collagen fiber coherency, and they enhanced the tensile strength of repaired rotator cuff tendon in an athymic rat model(11). AD MSCs decreased inflammatory cells and improved collagen fiber arrangement and organization as well as enhancing the tensile strength of rotator cuff tendon in a rat model of collagenase-induced rotator cuff injury(12). However, use of these MSCs has been criticized for several disadvantages including invasive techniques for harvesting(8), low collection efficiency(13), decreased quality with the age and morbidities of donors, and concerns of heterotopic ossification(14). Thus, it would be worthwhile to find alternate cell sources to overcome these drawbacks.

Umbilical cord derived MSCs (UC MSCs) are fetal MSCs isolated from umbilical cords which are usually discarded as medical waste after delivery, and thus can be obtained non-invasively at collection and can be supplied at relatively low cost(13). UC MSCs are reported to be more proliferative and have higher self-renewal potential than other adult MSCs(15). UC MSCs are able to differentiate towards the myogenic lineage and contributed to the muscle regenerative process of tibialis anterior muscle injury in a rat model(16). UC MSCs were also able to differentiate endothelial cells and contributed to the muscle regenerative process of a hindlimb ischemia injury in a mouse model(17). In addition, the conditioned medium of UC MSCs increased dermal fibroblast proliferation and migration and the medium accelerated wound healing with fewer scars of full-thickness skin excisional wounds injury in a C57BL6 mouse(18). Human umbilical cord perivascular cells facilitated tendon regeneration through changes in collagen organization, cell shape and orientation, and increase in mechanical properties in an athymic rat model of collagenase-induced Achilles tendon injury simulating a chronic tendon inflammation(19). Although few studies using UCB MSCs derived from fetal tissue have been reported for tendon healing (20, 21), but there is yet no report on the efficacy of UC MSCs for rotator cuff regeneration despite these potentials of UC MSCs for musculoskeletal tissue regeneration.

Hence, the purpose of this study was to investigate the efficacy of UC MSCs on the regeneration of a full-thickness rotator cuff defect (FTD) in a rat model. We hypothesized that UC MSCs would regenerate the FTD with tendon tissue similar properties to the normal tendon in terms of macroscopic, histological and biomechanical characteristics.

## Materials and Methods

### Study Design

This study was approved by the Institution’s Animal Care and Use Committee and the Institutional Ethical Review Board of our institute (IACUC_2017_0021 and IRB No. 16-2015-115). This study was carried out in strict accordance with the recommendations in the guide for the IACUC and IRB. All surgery was performed under anesthesia, and all efforts were made to minimize suffering. Eighty-four adult male Sprague-Dawley rats (12 weeks old, 340~360g) were divided into one of three groups and treated accordingly; 1) the control group (n=24); 2) the saline group (n=24); 3) the UC MSCs group (n=36). Rats from each group were sacrificed at three hours, two and four weeks after surgery. The supraspinatus tendon tissue (SST) was harvested and used for macroscopic (n=4; two and four weeks respectively) and histological evaluation (n=4; the tissue used for macroscopic evaluation is used for histology analysis), biomechanical evaluation (n=8; two and four weeks respectively), and cell trafficking evaluation (n=4; three hours, two and four weeks respectively).

### UC MSC Isolation and Culture

Human umbilical cords were obtained from healthy full-term deliveries by cesarean section after informed consent, and UC MSCs were isolated and cultured using minimal cube explant method.

UCs were washed 2-3 times with Dulbecco’s Phosphate-Buffered Saline (DPBS; Welgene, Daegu, Korea) to remove blood products and then measured for length and weight, and then cut into minimal cube explants of 2-4 mm by surgical scissors. The cube explants (1 g) were aligned at regular intervals in 15-cm culture dishes and allowed to firmly attach to the bottom of the dish for 60 mins in a 5 % CO_2_ incubator with humidified air at 37 °C. Then, the culture medium consisting of low-glucose Dulbecco’s modified Eagle medium (LG-DMEM; Hyclone, Logan, USA) supplemented with 10 % fetal bovine serum (FBS; Hyclone) and antibiotic-antimycotic solution (100 U/ml penicillin, 100 μg/ml streptomycin, and 0.25 μg/ml amphotericin B; Welgene) was gently poured into the dishes. The medium was replaced twice a week. Non-adherent cells were removed by medium changes. When cells reached 60–80% confluence, they were detached by incubation for 3 min with trypsin-EDTA (0.05% trypsin, 0.53 mM EDTA; Welgene). The tissues were removed through a 100 μm cell strainer (SPL Life Sciences, Pocheon, Korea) and the cells were centrifuged at 500 g for 5 min at 20 °C and then replated at a density of 3 x 10^3^ cells/cm^2^.

Surface phenotypic characterization by flow cytometry was performed as previously reported (22). Briefly, a total of six antibodies were used: CD34-PE, CD45-PE, CD73-PE, CD90-FITC, CD105-FITC and HLA-DR-FITC (BD Biosciences, San Jose, CA, USA). Once the cells had been detached, aliquots of 5×10^5^ cells were washed twice with DPBS, centrifuged, washed in ice cold DPBS supplemented with 2% fetal bovine serum (FCM buffer) and fixed in 1% paraformaldehyde in FCM buffer, followed by incubation with fluorescein isothiocyanate (FITC) or phycoerythrin (PE)-conjugated antibodies for 30 min on 4 °C in a dark room. Data were obtained by analyzing 10,000 events on a Flow Cytometer FC500 (Beckman Coulter, Brea, CA, USA) with CXP Cytometer 2.3 software (Beckman Coulter, Brea, CA, USA). All analyses were standardized against isotype controls.

UC MSCs at passage 10 were used for all experiments and characterization including morphology, growth kinetics, CFU-F, flow-cytometric and trilineage differentiation of cells was performed as previously reported (22).

### Surgical Procedures

Anesthesia was induced using zoletil and rompun (30mg/kg + 10mg/kg). The left shoulder was operated on in all cases. A 2cm skin incision was made directly over the anterolateral border of the acromion. After the SST was exposed by detaching trapezius and deltoid muscle from the acromion, a round FTD with a diameter of 2mm in the middle of the SST was created 1mm from the insertion using a Biopsy Punch (BP-20F, Kai Medical Europe GmbH, Bremen, Germany). This defect size was approximately 50% of the tendon width, correlating to a large but not a massive tear according to the method previously described(23, 24). Ten microliter of saline, UC MSCs (1 x 10^6^ cells) and PKH26 labeled UC MSCs (1 x 10^6^ cells) were intratendinously injected adjacent to both sides of the defect in two divided doses via 30 G needle (Fig 1A). After injection, the deltoid and trapezius muscles were sutured with a 4-0 Vicryl suture (W9074, Ethicon, Cincinnati, OH, USA) and then skin was also sutured with a black silk (SK439, AILee, Busa, Korea). In the control group, the SST was exposed, but no further surgical procedure was performed (the sham control). After surgery, animals were allowed free cage activity. The condition of animals were checked by veterinarians and zookeepers.

**Fig 1.**
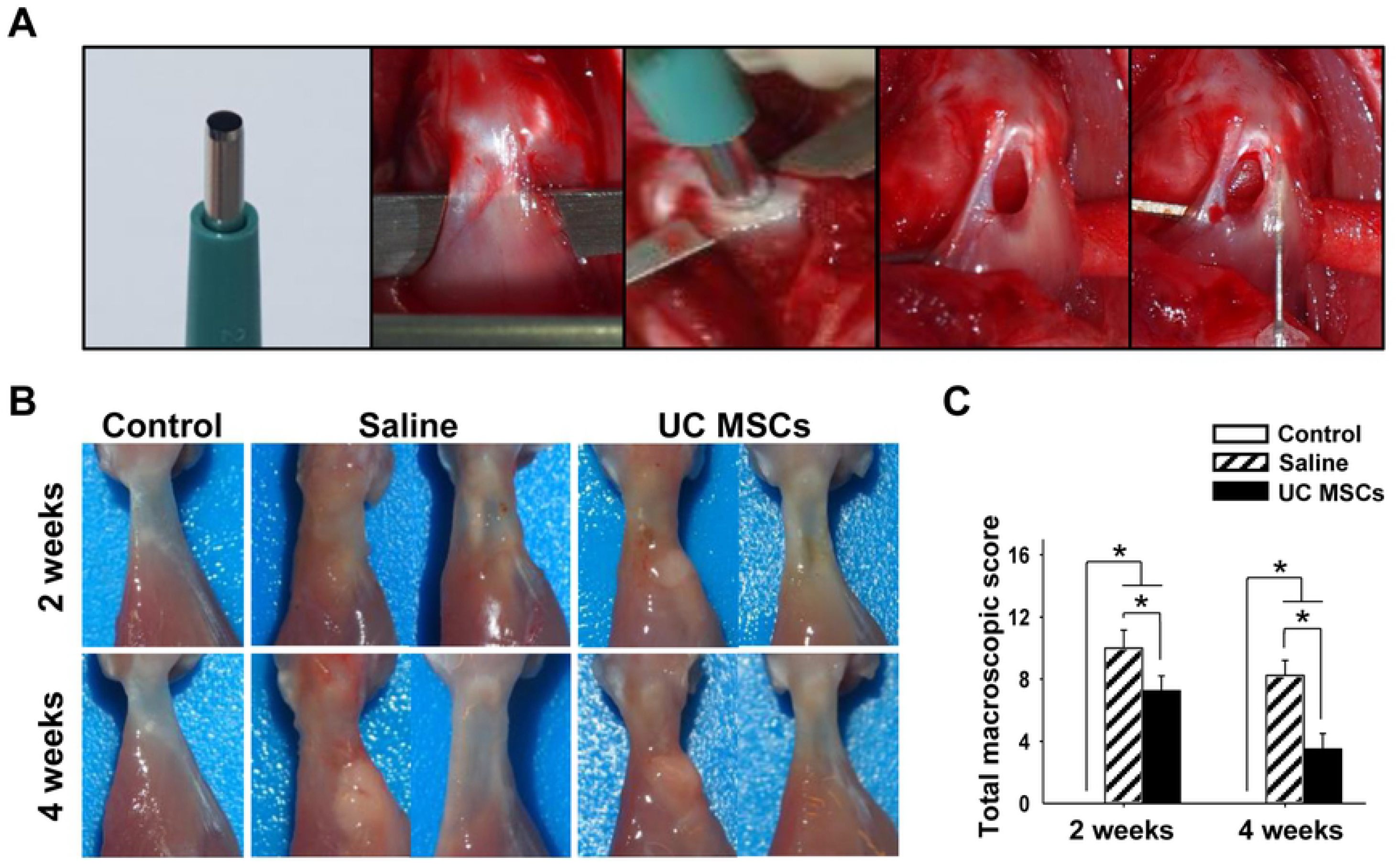
Procedure of Surgery, macroscopic images, and quantification of macroscopic appearance of regenerated tendons at two and four weeks. (A): Surgical procedure of FTD and intratendinous injection of saline or UC MSCs. (B): Macroscopic appearance of the tendon (Left side image: Supraspinatus tendon immediately after harvest; Right side image: the tendon removing the loose connective tissue surrounding the injury site to observe the original defect shape. (C): The total macroscopic score. Bar charts present mean ± standard deviation; Statistically significant at *p* < .05. Abbreviations: UC MSCs, umbilical cord derived mesenchymal stem cells.

### Macroscopic Evaluation

At two and four weeks after injection, the rats were sacrificed in a carbon dioxide chamber. The SST of the rats were harvested with humerus head without removing the muscle. For macroscopic evaluation of tendon regeneration, we used a modified semi-quantitative system of Stoll *et al.(25)* (see the S1 table). The total macroscopic score could vary between 0 (normal tendon) and 15 (most severely injured).

### Histological Evaluation

After the macroscopic evaluation, the harvested tissues were immediately fixed in 4% (w/v) paraformaldehyde (PFA; Merck, Darmstadt, Germany) for 24 hours, and underwent decalcification in 10% ethylendiaminetetracetic acid (EDTA; Sigma-Aldrich, St Louis, MO, USA) for two days. After decalcification, the tissues were dehydrated through an increasing ethanol gradient, defatted in chloroform, and embedded in paraffin blocks. The tissue was carefully trimmed to the appropriate middle site of tendon and cut into 4-μm thick serial sections.

A randomly selected slide was stained with hematoxylin and eosin (H&E) and analyzed by light microscopy (U-TVO 63XC; Olympus Corp., Tokyo, Japan). For the evaluation of tendinopathy, each slide was evaluated using the semi-quantitative grading scale as previously described (26). The 7 parameters of the system include; fiber structure, fiber arrangement, rounding of the nuclei, variations in cellularity, vascularity, stainability, and hyalinization. Each parameter in the grading scales varies from 0 to 3. The total degeneration score for a given slide varied between 0 (normal tendon) and 21 (most severely degenerated).

Integration of structure and infiltration of inflammatory cells were evaluated using a 0-3 grading scale: 0 (normal), 1 (slightly abnormal), 2 (moderately abnormal) and 3 (maximally abnormal) (27).

We also evaluated the occurrence of heterotopic ossification when separated, clustered and bar-shaped foci were found in the whole tendon structure (28).

In normal tendon, the few fibroblasts with flattened nuclei are typically aligned parallel to the tensile axis. After injury, the morphometric changes of fibroblast nuclei were evaluated as previously described by Fernandez-Sarmiento using H&E stained slides (29). Fibroblast density (number of nuclei per mm^2^), nuclear aspect ratio (the ratio of the minor diameter to the maximal diameter), and nuclear orientation angle (between the major axis of the nuclear angle and the axis of collagen fibers) were evaluated. Five regions of interest (ROI) were measured and the average was used finally.

Slides were stained with picrosirius red (PSR) for analysis of collagen fiber organization and coherency using circularly polarized light microscopy at x200 magnification. Collagen organization was measured as intense white areas of brightly diffracted light on gray scale (black, 0; white, 255) using ImageJ software with installed NII plugin (National Institutes of Health, MD, USA). Higher gray scale indicated more organized and mature collagen (30). The coherence of the collagen fibers is a measure of the extent of fiber alignment in the major axis of alignment. The coherence was quantified using the Orientation J plug-in for image J and then multiplied by 100 to obtain the final coherence value (11). Five ROIs were measured and the mean value was used.

For evaluation of cartilage formation, slides stained with Safranin-O/fast green (Saf-O) were used and observed via light microscopy at x 200 magnification. The glycoaminoglycan (GAG)-rich area was measured using image J (14).

### Biomechanical Evaluation

For biomechanical testing, we harvested supraspinatus tendon with humerus head and carefully removed the muscles to leave only the tendon. The harvested tissues were wrapped in saline-soaked gauze and kept at −80 °C. Before testing, the tissues were thawed with saline wet gauze at room temperature for 24 hours and the tissues were kept moist with saline during all tests. The distal part of humerus bone was vertically embedded in an aluminum tube full of polymethylmethacrylate (PMMA) in the custom-designed lower jig of a testing system. The proximal end of tendon was compressed with sandpaper, gauge, and rubber to prevent slippage and to reduce damage of specimens. The complex was clamped vertically in the custom-designed upper jig. Testing was performed with shoulders at 90° of abduction with a material testing system (H5K5, Tinus Olsen, England, UK) (31, 32). All specimens were loaded to failure in tension at a constant rate of 0.1 mm/s. Slippage of the tendon was inspected visually. The cross-sectional area of supraspinatus tendon was measured at the center of defect region. The area was calculated with the formula (Area = width x thickness x π). From the load-displacement curve recorded during tests, the ultimate failure load, the stiffness and ultimate stress was calculated (33).

### UC MSCs Trafficking

UC MSCs were labeled with fluorescent PKH26 (Sigma-Aldrich, St Louis, MO, USA) according to the manufacturer’s protocol that is known to have an *in vivo* half-life of greater than 100 days and would be useful for *in vivo* cell trafficking(34). When cells reached 60% to 80% confluence, they were detached by incubation for 5 minutes with 0.25% trypsin EDTA and washed three times with PBS After aspirating the supernatant in UC MSC containing tubes, the cells were resuspended gently in 1mL of the dilution buffer and mixed with an equal volume of the labeling solution containing 4×10^-2^_M_ PKH26 in the dilution buffer. Then, the cells were incubated at 5 min at room temperature and the reaction was terminated by addition 2 mL of FBS. The suspension was centrifuged at 400xg for 5 min and the cells were washed three times with PBS. After labeling, the cells were counted by hemocytometer, and confirmation of red fluorescence was obtained by fluorescence microscope (Leica DMI 4000B, Leica, Wetzlar, Germany) before injection.

At three hours, two and four weeks after injection, the tissue was harvested and used for evaluation. We harvested only supraspinatus tendon and the harvested tissues were immediately fixed in 4% (w/v) PFA for 24 hours. The tissues were treated with consecutive 10%, 15% and 20% sucrose/PBS solutions at 4 °C for 12h respectively. The tissues were embedded in O.C.T. compound (Tissue-Tek, Miles, USA) with subsequent freezing of the block at −80 G. To evaluate the survival ability of the PKH 25 labeled UC MSCs, the specimens were sectioned (7 μm) with a freezing microtome (Leica CM3050S, Leica, Wetzlar, Germany) and carefully trimmed until we found middle side of tendon. The slides were mounted with Vectashield containing DAPI (Vector Laboratories Inc., Burlingame, CA). Five fields were randomly selected in the slide and high-powered images (x400) were obtained by fluorescence microscopy. The PKH26 positive cells coincident with 4’,6-diamidino-2-phenylindole (DAPI) were counted per area and the mean number were recorded by image J(35).

### Statistical Analysis

All data are shown as mean ± SD. The data was analyzed with one-way analysis of variance (ANOVA) with post hoc analysis using Bonferroni multiple comparison test. All statistical analyses were performed with SPSS software version 23 (IBM). Differences of *p* < .05 were considered statistically significant.

## Results

### Macroscopic Evaluation

For macroscopic scoring, tendon thickness score was significantly less in the UC MSCs group in comparison with that in the saline group ant the total macroscopic score was significantly lower in the UC MSCs group, 7.25 ± 0.96, than in the saline group, 10.00 ± 1.15 *(p* = .005) at two weeks. At four weeks, inflammation, defect size, swelling/redness and connection surrounding tissue and slidability score were significantly lower in the UC MSCs group than those in the saline group. The total macroscopic score was significantly lower in the UC MSCs group, 3.50 ± 1.00, than in the saline group, 8.25 ± 0.96 (*p* < .001) at four weeks (Figs 1B and 1C). Loose connective tissues surrounding the defect site was removed to observe the original defect (Fig 1B).

### Histological Evaluation

For the evaluation of tendinopathy, stainability score was significantly less in the UC MSCs group, 1.50 ± 0.58, in comparison with that in the saline group, 2.50 ± 0.58 (p = .045) at two weeks. However, the total degeneration score was not significantly different between the saline group, 16.50 ± 1.00, and the UC MSCs group, 15.50 ± 1.73 (p = .905) at two weeks (Figs 2A and 2B). At four weeks, fiber structure, fiber arrangement, rounding of the nuclei vascularity and hyalinization were significantly lower in the UC MSCs group, 1.25 ± 0.50, 1.00 ± 0.00, 1.00 ± 0.00, 1.25 ± 0.50 and 0.25 ± 0.50 than those in the saline group, 2.50 ± 0.58, 2.50 ± 0.58, 2.25 ± 0.50, 2.50 ± 0.58 and 1.75 ± 0.50 *(p* = .009, < .001, .006, .025 and .002 respectively). The total degeneration score was significantly less in the UC MSCs group, 7.00 ± 1.41, than in the saline group, 15.67 ± 2.08 *(p* < .001) (Figs 2A and 2B) at four weeks. The scores of structure integration and inflammation were not significantly different between the saline and UC MSCs groups at both two and four weeks (Figs 2A, 2C and 2D). There was no noticeable immune response against the UC MSCs. Heterotopic ossification was not observed in both groups at any time points.

**Fig 2.**
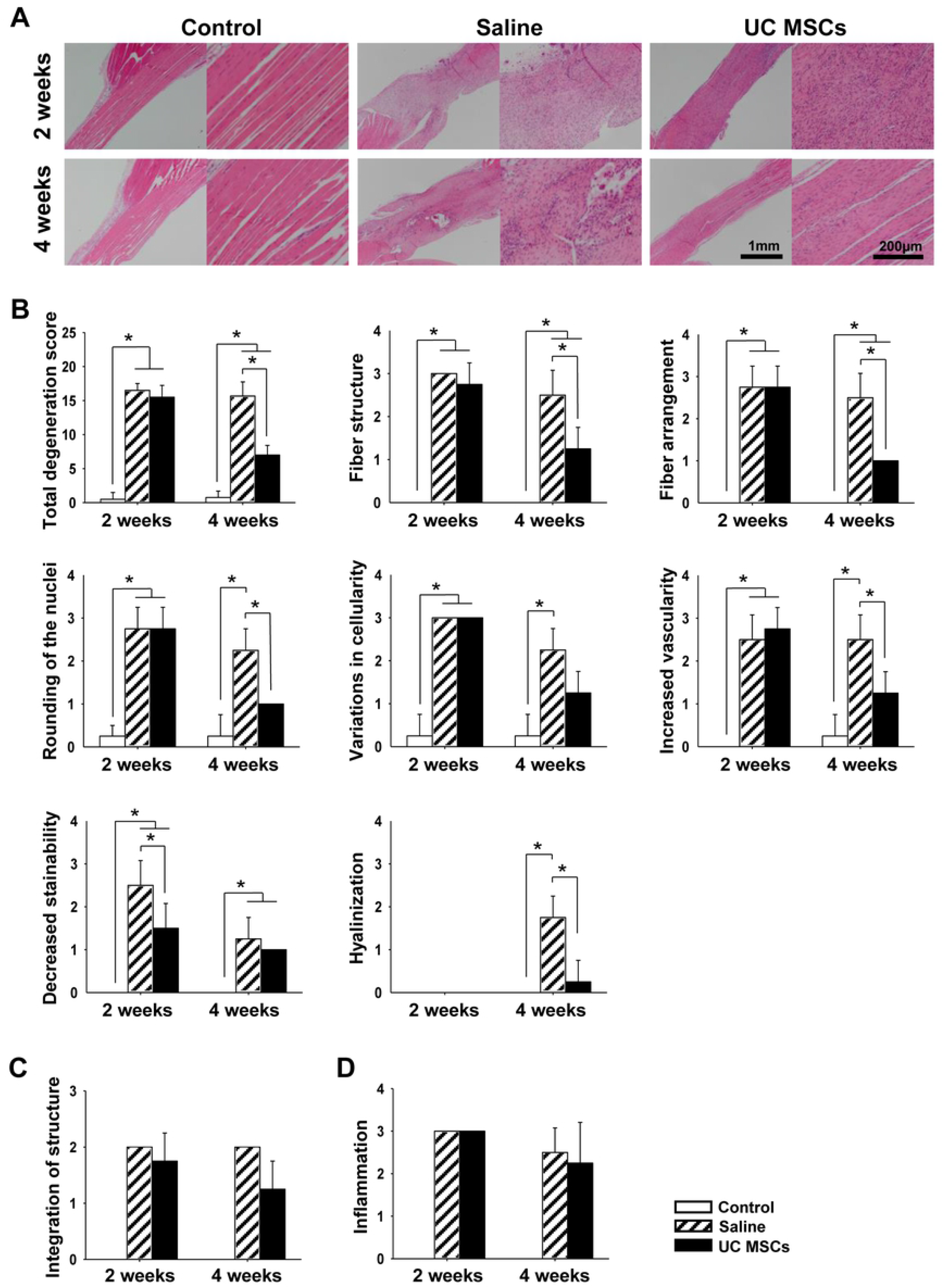
Representative histological images of general structure and quantification of the changes in regenerated tendon at two and four weeks. (A): H&E staining of tendon (magnification; X40 and X200). (B): the total degeneration score and detail parameters. (C): Integration of structure between the tendon defect and adjacent tendon. (D): Inflammation at the tendon defect. Bar charts present mean ± standard deviation; statistically significant at *p* < .05. Abbreviations: UC MSCs, umbilical cord derived mesenchymal; H&E, hematoxylin and eosin

Collagen organization score was not different between the groups at two weeks, but it became significantly higher in the UC MSCs group, 108.79 ± 12.36, than in the saline group, 78.67 ± 8.71 *(p* = .005) at four weeks (Fig 3A and 3B). Collagen fiber coherence score was not different between the groups at any time points (Figs 3A and 3C).

**Fig 3.**
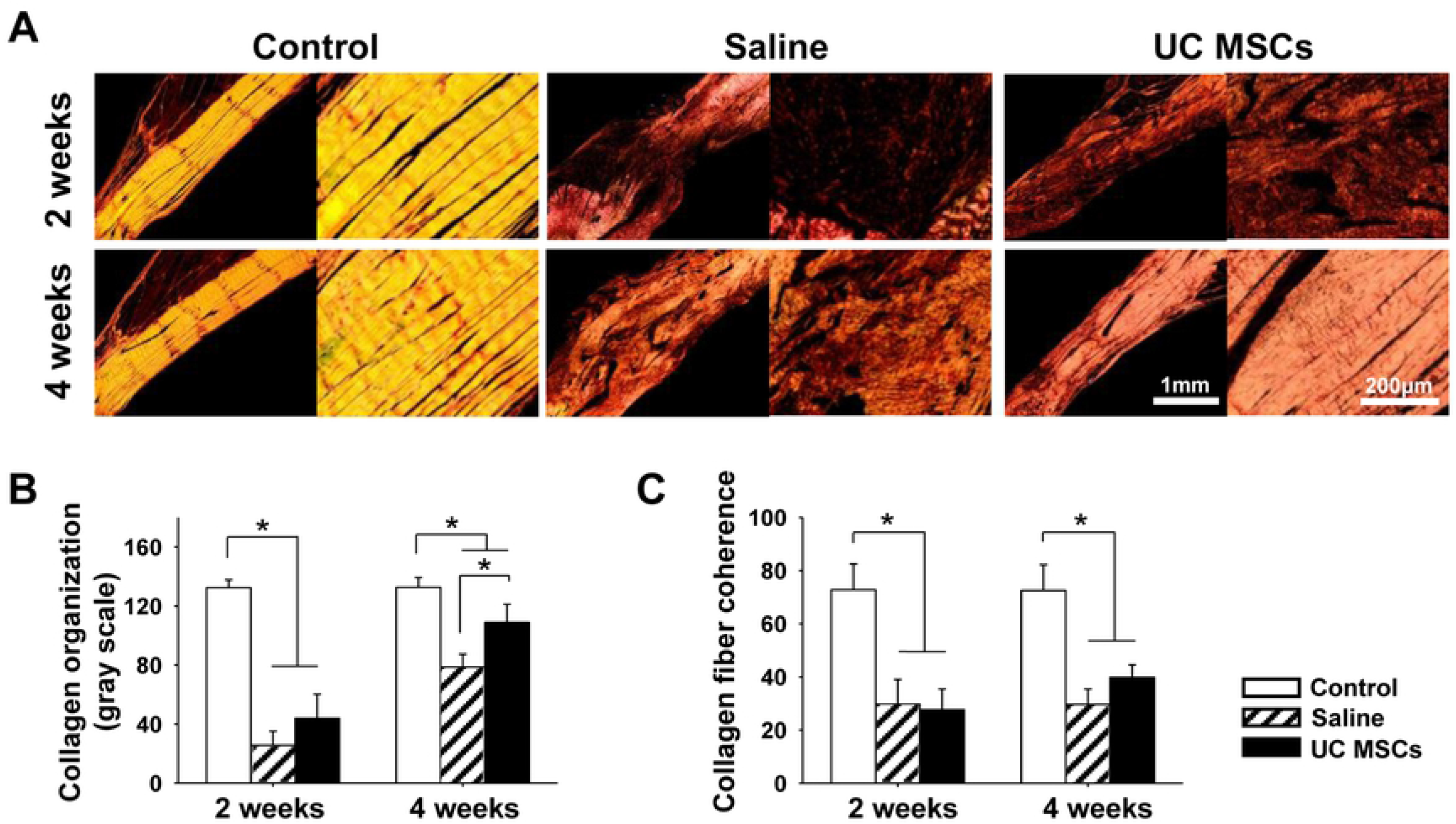
Representative histologic images of collagen matrix and quantification of the changes in regenerated tendon at two and four weeks. (A): PSR staining of the tendon (magnification; X40 and X200). (B): collagen organization of the tendon. (C): collagen fiber coherency of the tendon. Bar charts present mean ± standard deviation; statistically significant at *p* < .05. Abbreviations: UC MSCs, umbilical cord derived mesenchymal; PSR, picrosirius red.

Fibroblast density was not significantly different between the groups at any time points (Figs 4A and 4B). Scores of nuclear aspect ratio and nuclear orientation angle were not significantly different between the two groups at two weeks. At four weeks, they became significantly lower in the UC MSCs group, 0.21 ± 0.02 and 6.49 ± 1.52, than in the saline group, 0.35 ± 0.07 and 21.61 ± 7.76 respectively *(p* = .004 and .004) at four weeks (Figs 4A, 4C and 4D).

**Fig 4.**
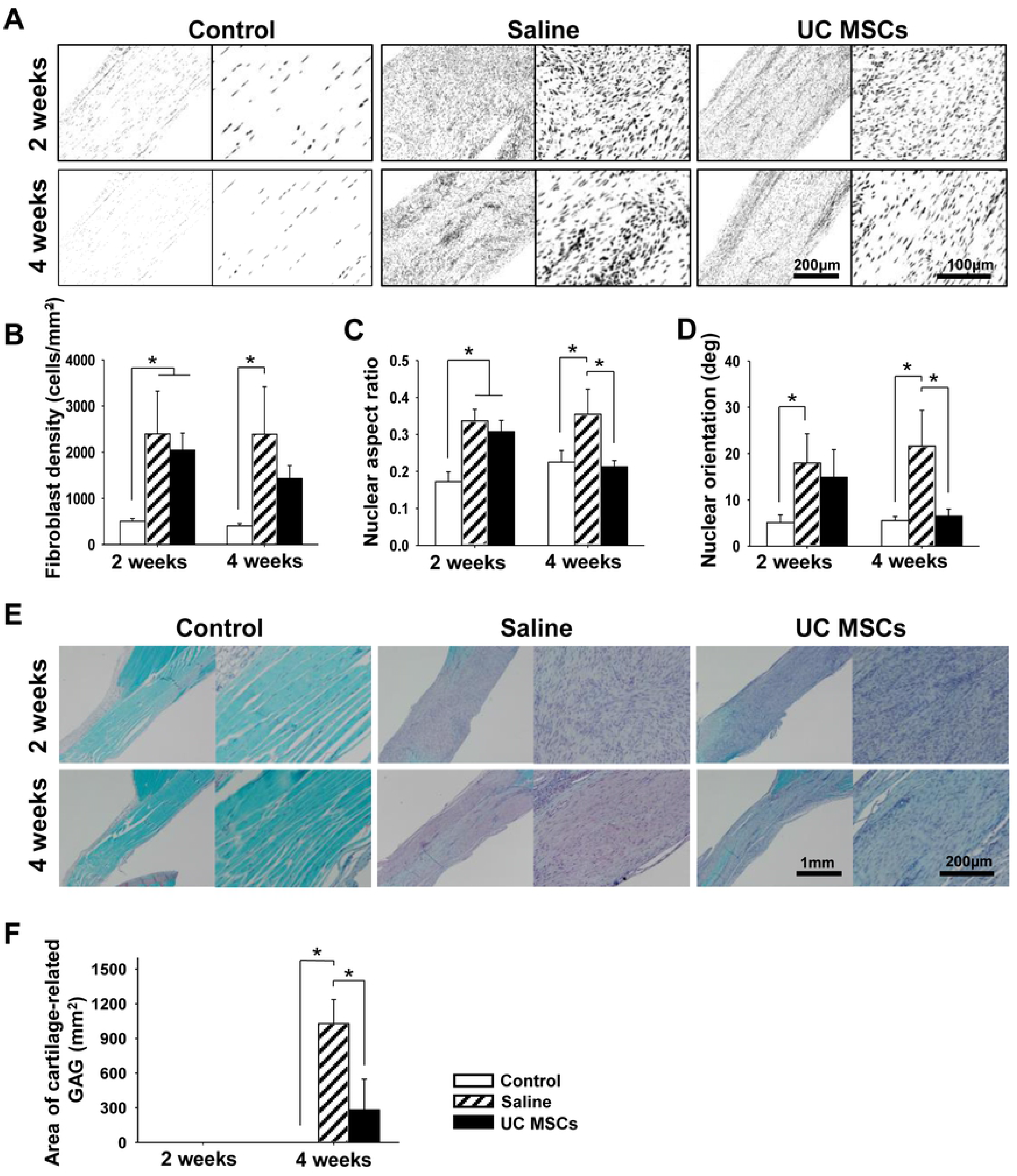
Representative histological images of nuclei morphology and cartilage formation of regenerated tendons and quantification of the changes at two and four weeks. (A): Nuclei morphology of fibroblasts at the tendon (magnification; X200 and X400) (B): Density of fibroblasts of the tendon. (C): Nuclear aspect ratio of fibroblasts of the tendon. (D): Nuclear angle of fibroblasts of the tendon. (E): Saf-O staining of the tendon (magnification; X40 and X200). (F): Area of cartilage-related GAG of the tendon. Bar charts present mean ± standard deviation; statistically significant at *p* < .05. Abbreviations: UC MSCs, umbilical cord derived mesenchymal; GAG, glycoaminoglycan; Saf-O, safranin-O.

Area of cartilage-related GAG was not found in either group at two weeks. At four weeks, the GAG-stained area appeared in both the saline and the UC MSCs groups, and it was significantly less in the UC MSs group, 281.01 ± 267.23 mm^2^, than that in the saline group, 1,030.59 ± 206.12 mm^2^, *(p* = .001) (Figs 4E and 4F).

### Biomechanical Evaluation

The cross-sectional area of the SST significantly increased in the saline and the UC MSCs groups, 3.12 ± 0.14 and 3.02 ± 0.24 mm^2^, in comparison with that in the control group, 2.29 ± 0.22 mm^2^, at two weeks *(p* < .001 and < .001) (Fig 5B). At four weeks, the cross-sectional area was significantly less in the UC MSCs group, 2.84 ± 0.23 mm^2^, than that in the saline group, 3.28 ± 0.64 mm^2^ *(p* < .001).

**Fig 5.**
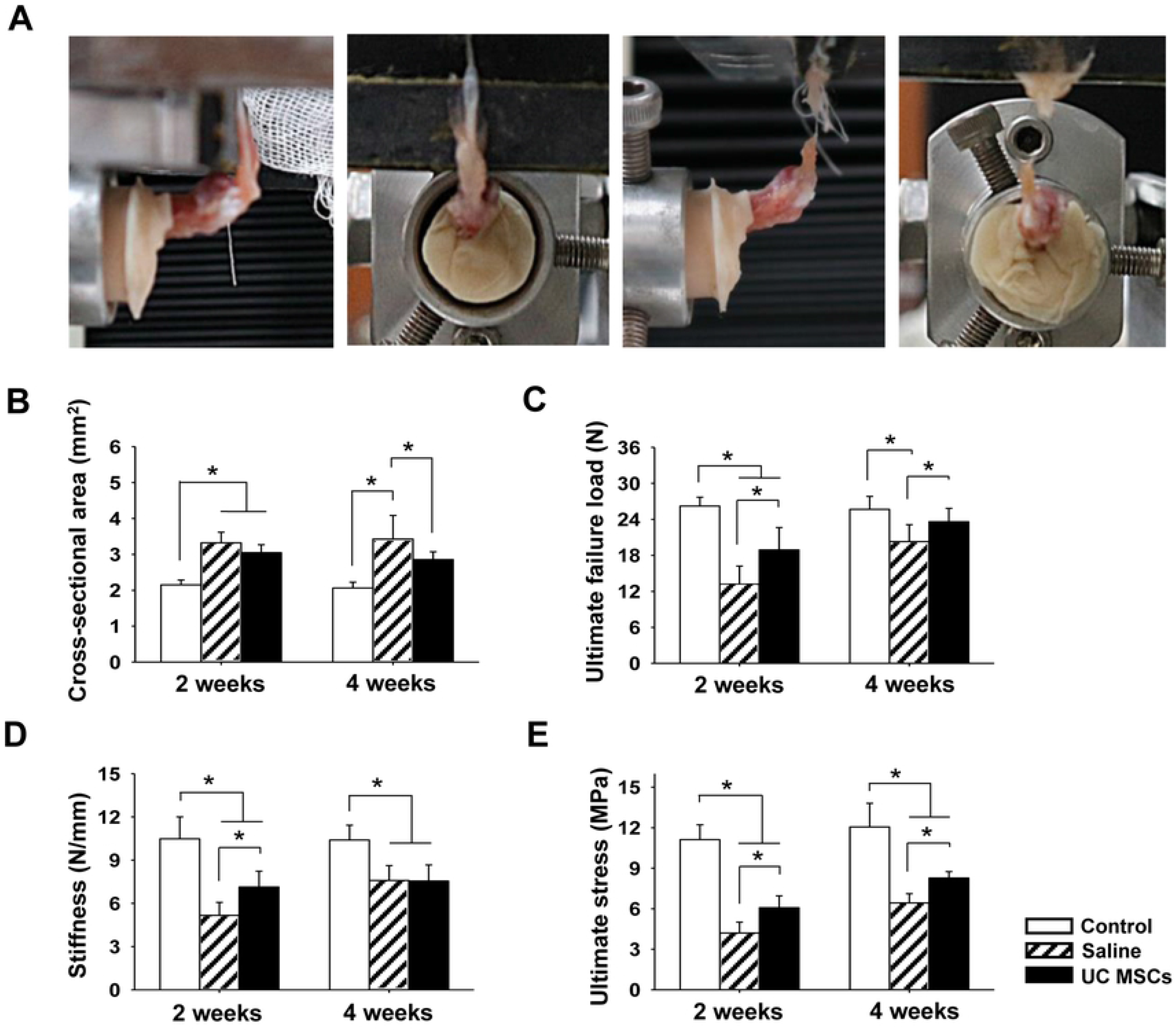
Biomechanical test procedure and quantification of biomechanical properties of regenerated tendons at two and four weeks. (A): a extracted specimen and procedure of biomechanical testing. (B): Cross-sectional area of the supraspinatus tendon at defect site. (C): ultimate failure load. (D): stiffness. (E): ultimate stress. Bar charts present mean ± standard deviation; statistically significant at *p* < .05. Abbreviations: UC MSCs, umbilical cord derived mesenchymal stem cells.

The ultimate failure load was significantly higher in the UC MSCs group, 17.58 ± 3.52 N and 22.68 ± 2.44 N, than those in the saline group, 13.18 ± 3.04 N and 18.37 ± 4.31 N, at two and four weeks, respectively *(p* = .016 and .035) (Fig 5C). The ultimate failure load in the UC MSC group, 22.68 ± 2.44 N, at four weeks was comparable with that of the control group, 25.67 ± 2.17 N *(p* = .159).

The stiffness significantly improved in the UC MSCs group, 6.99 ± 1.05 N/mm than that in the saline group, 5.18 ± 0.90 N/mm at two weeks *(p* = .018) (Fig 5D). However, it was not different between the two groups at 4 weeks.

The ultimate stresses were significantly higher in the UC MSCs group, 5.77 ± 0.79 MPa and 7.97 ± 0.52 MPa, than those in the saline group, 4.20 ± 0.81 MPa and 5.84 ± 1.68 MPa, at two and four weeks, respectively *(p* = .007 and *p* < .001) (Fig 5E). However, the ultimate stress in both groups at four weeks was yet significantly less than that in the control group, 11.99 ± 1.89 MPa *(p* < .001 and < .001, respectively).

### UC MSCs Trafficking

Intratendinously injected UC MSCs disappeared over time, but they were retained in the tendon defect until 4 weeks after injection. At three hours after UC MSCs injection, the number of PKH 26 positive cells was 2135.36 ± 437.02 cells/mm^2^ (Figs 6A and 6B). At two and four weeks, it significantly decreased to 454.59 ± 137.40 cells/mm^2^, 21.29% of that at three hours, and to 113.01 ± 21.39 cells/mm^2^, 5.29% *(p* < .001 and <.001, respectively).

**Fig 6.**
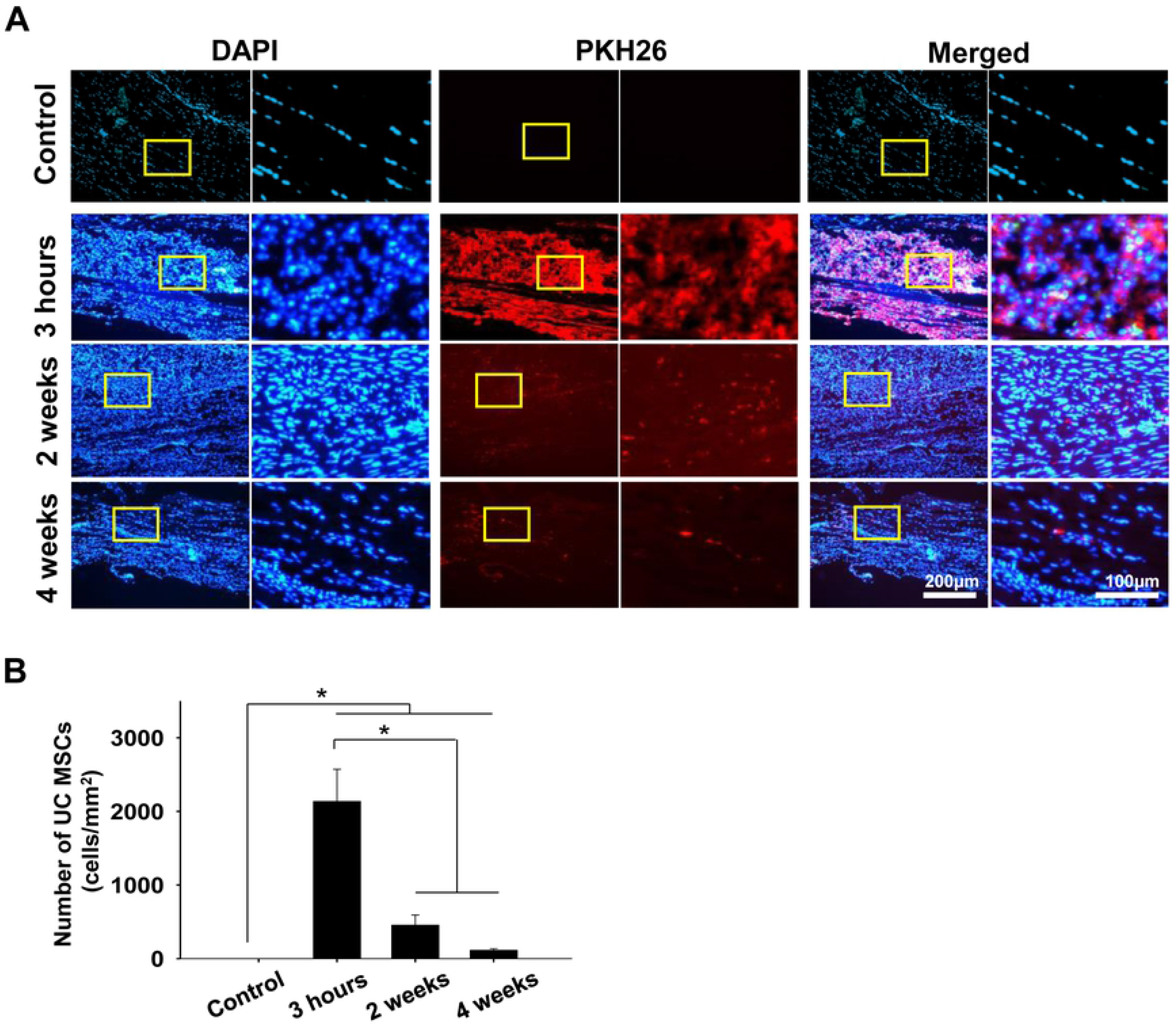
Representative fluorescent images of implanted PKH26 labeled UC MSCs within the SST and quantification of the injected cells at 3 hours and two and four weeks. (A): Implanted UC MSCs with DAPI and PKH26 dye of the tendon (magnification; X200 and X400). (B): The number of the injected UC MSCs per area (mm^2^). Bar charts present mean ± standard deviation; statistically significant at *p* < .05. Abbreviations: DAPI, 4’,6-diamidino-2-phenylindole; UC MSCs, umbilical cord derived mesenchymal stem cells.

## Discussion

The most important findings of this study are as follows; 1) the total macroscopic score significantly improved in the UC MSCs group compared to the saline group at two and four weeks through tendon thickness, inflammation, defect size, swelling/redness and connection surrounding tissue and slidability; 2) histologically, the total degeneration score significantly improved in terms of fiber structure, fiber arrangement, rounding of the nuclei, vascularity and hyalinization as well as the scores of nuclear aspect ratio, nuclear orientation angle, and collagen organization in the UC MSCs group than in the saline group at four weeks; 3) the area of cartilage-related GAG was significantly less in the UC MSCs group than in the saline group at four weeks and heterotopic ossification was not observed at any groups; 4) the ultimate failure load was significantly greater in the UC MSCs group than that in the saline group at two and four weeks. Especially, the ultimate failure load in the UC MSCs group at four weeks was comparable with that of the control group; and 5) intratendinous injected UC MSCs disappeared over time, and 5.29% of those at three hours were retained in the tendon defect at four weeks after injection. Taken together, these results showed that intratendinous injection of UC MSCs would affect regeneration of the FTD with the formation of collagen matrix while suppressing heterotopic matrix changes, and thus improved the ultimate failure load comparable to normal tendon at 4 weeks in a rat model suggesting potentials of UC MSCs injection for the treatment of the FTD without surgical repair.

Clinically, rotator cuff disease is known to progress from tendinopathy to the PTD, and then to the FTD(26). The FTD has been reported to increase in size over time(36). Progression of rotator cuff disease does not only mean an increase of the size of the FTD, but also the deterioration of tendon structure and quality, and degeneration of rotator cuff muscles(37, 38). Therefore, it would be crucial to prevent or halt, or should be better to reverse its progression though the regeneration of the FTD. Several researchers have shown regenerative potentials of MSCs in patellar tendon defect model, Achilles tendon defect and incision models, superficial digital flexor tendon core lesion model and Achilles tendinopathy model(39, 40). For surgical repair model of rotator cuff tear, several studies demonstrated that MSCs from BM, AD, and tendon implanted with scaffold have shown promising results in repair and defect models(41, 42). However, there were few studies that delivered MSCs via injection for the investigation of regeneration of rotator cuff tendon in animal model; one study injected UCB MSCs with hyaluronic acid into subscapularis defect in a rabbit model, and one injected AD MSCs in phosphate buffered saline into tendinopathy lesion of SST in a rat model(12, 43). In this study, we injected human UC MSCs adjacent to the FTD in a rat model without any carrier which could be feasibly translated in daily practice. In this study, we observed UC MSCs could prevent rotator cuff tear from progression to worst over time by preventing the tear size from increasing, whereas the defect size increased in the saline group. Furthermore, tendon thickness, inflammation, defect size, swelling/redness and connection surrounding tissue and slidability also improved in the UC MSCs group compared to the saline group. These results demonstrated healing potentials of injected UC MSCs without carrier for the treatment of the FTD which is consistent with that of the previous study.

The main cause of unsuccessful tendon healing is the formation of a fibrovascular scar tissue after injury(44). Scar tissue has decreased and disorganized collagen matrix with reduced collagen 1/3 ratio(45), as well as increased vascularity and angled and rounded nuclei of fibroblasts(46). Previously, there are several studies trying to return the degenerated tendon structure to its previous normal tendon structure using MSC treatments(47, 48). BM MSCs improved general tendon structure such as fiber structure, arrangement, in Achilles tendon rupture rat model at 30 days(48) and AD MSCs improved matrix organization in Achilles tendon defect model at 3 and 4.5 weeks(47). Nonetheless, these results did not clearly show that BM and AD MSCs reduced scar tissue formation, because histological criteria for the scar tissue has not been clear(47, 48). In this study, we established a histological evaluation system that encompassed various currently accepted methods that could define scar tissue in tendon. General tendon structure including collagen matrix, cell, vascularity, hyalinization, integration and inflammation was evaluated by H&E staining and light microscopy, and collagen fiber organization and coherency were analyzed by PSR staining and polarized light microscopy. Heterotopic matrix changes were evaluated by Saf-O staining using light microscopy. Furthermore, we used computerized image analysis to quantification of changes of tendon structure. With this system, we found UC MSCs could improve tendon structure suppressing abnormal scar tissue formation and the regenerated tendon structure was comparable to normal tendon tissue.

Cartilage-related GAG rich area or hyalinization, calcific tendinopathy and heterotopic ossification has been commonly reported in patients with tendinopathy and tendon rupture(49). In addition, chondrocyte phenotype and ectopic bone formation has been frequently described in surgically induced defect models and collagenase-induced tendinopathy models(50, 51). Furthermore, BM MSCs application for tendon regeneration evoked heterotopic ossification both in defect and tendinopathy models(14, 52). The presence of heterotopic ossification in tendon worsens the clinical manifestations of tendon injury(53), may lead to increase rupture rates and recovery time(54), and cause a higher frequency of post-operative complications such as shoulder pain(55). Therefore, it is crucial to suppress heterotopic ossification for the treatment of tendon injury with MSC implantation. With respect to the occurrence of heterotopic ossification, current evidences raise concerns over the use of BM MSCs for tendon regeneration because they were prone to osteogenic differentiation with a higher chance of heterotopic ossification up to 55% of cases in experimental studies. However, in this study, UC MSCs reduced heterotopic cartilage-related GAG formation by 73% of saline treated tendons at four weeks and did not cause any heterotopic ossification. Although it had not been studied how to inhibit heterotopic cartilage or bone formation by UC MSCs, these results demonstrated that UC MSCs could suppress heterotopic matrix changes and ossification suggesting that UC MSCs is a safer option than BM MSCs for tendon regeneration.

Several studies showed that MSCs application improved some of biomechanical properties of surgically repaired and collagenase-induced tendinopathy models(12, 39). However, the enhanced properties provided with MSCs may not last longer time or some parameters such as ultimate stress did not improve with application of MSCs(12, 39, 47). In this study, UC MSCs injection significantly increased the ultimate failure load and ultimate stress at two and four weeks, and also improved stiffness at two weeks compared to the saline injection. Especially the ultimate failure load increased by 88% to a level comparable to the normal tendon at four weeks. Furthermore, the ultimate stress in the UC MSCs group were significantly higher by 1.4-fold both at two and four weeks after injection than that in the saline group. These improved biomechanical results were corroborated by the improved histologic results in this study. Biomechanical properties of tendon are derived from well aligned and organized collagen fibers of tendon (47, 56). Thus, we tough that UC MSCs could promote the recovery of the tendon structure, then the recovered tendon structure could induce the functional recovery of the rotator cuff.

There are several limitations in this study. First, an acute tendon defect model which might not reflect chronic tendon degeneration, one of the most important causes of rotator cuff tear, was used. However, the total degeneration scores in the saline group at two and four weeks were similar to those in patients with a full-thickness Second, evaluation period of four weeks would be short for clearer assessment of effects of UC MSCs for rotator cuff tendon regeneration as lots of parameters in the study were expected to further improve in a longer follow-up period.

This study demonstrated that UC MSCs regenerated the FTD with tendon tissue similar properties to the normal tendon in terms of macroscopic, histological and biomechanical characteristics in a rat model through directly inject to the injury site without any carrier materials.

## Acknowledgments

This research was supported by a grant (NRF-2015M3A9E6028412) of the Bio & Medical Technology Development Program and a grant (NRF-2017R1A2B2010995) of the Basic Science Research Program of the National Research Foundation of Korea. C.H.J. owns shares of AcesoStem Biostrategies Inc. The other authors indicated no potential conflicts of interest.

## Authors’ contributions

Conceptualization: J.H.Y., C.H.J.

Data curation: J.H.Y., T.S.B.

Formal analysis: J.H.Y.

Investigation: J.H.Y.

Methodology: J.H.Y.

Visualization: J.H.Y.

Writing-original draft preparation: J.H.Y.

Resources: J.H.Y., I.J.K., G.Y.S., J.K.P., A.Y.L., B.C.C., T.S.B., and B.J.K.

Funding acquisition: C.H.J.

Project administration: C.H.J.,

Supervision: C.H.J.

Writing-review & editing: T.S.B., C.H.J.

## Supporting information

S1 Table. Modified macroscopic evaluation system table.

